# Prevalence, awareness and control of hypertension in Malaysia 1980 – 2017: A Systematic Review and Meta-Analysis

**DOI:** 10.1101/625004

**Authors:** Zhen Yee Chow, Soo Man Jun, Siew Mooi Ching, Chun Han Tan, Kai Wei Lee, Navin Kumar Devaraj, Hani Syahida, Vasudevan Ramachandran, Fan Kee Hoo, Ai Theng Cheong, Yook Chin Chia

**Affiliations:** Faculty of Medicine and Health Sciences, University Putra Malaysia, 43400 Serdang, Malaysia; Postgraduate School, Faculty of Medicine and Health Sciences, University Putra Malaysia, 43400 Serdang, Malaysia; Department of Family Medicine, Faculty of Medicine and Health Sciences, University Putra Malaysia, 43400 Serdang, Malaysia; Malaysian Research Institute on Ageing, Universiti Putra Malaysia, 43400 Serdang, Malaysia; Department of Medicine, Faculty of Medicine and Health Sciences, University Putra Malaysia, 43400 Serdang, Malaysia; Department of Medical Sciences, School of Healthcare and Medical Sciences, Sunway University, Bandar Sunway, 47500 Selangor, Malaysia; Department of Primary Care Medicine, Faculty of Medicine, University of Malaya, Kuala Lumpur, Malaysia

**Keywords:** Prevalence, Awareness, Control, Hypertension, Blood Pressure, Malaysia, Systematic review, Meta-analysis

## Abstract

**Background:** Hypertension is a common public health problem worldwide and is a well-known risk factor for increased risk of cardiovascular diseases, contributing to high morbidity and mortality. However, there is no systematic review and meta-analysis that has been done in a multi-ethnic population like Malaysia. This systematic review aims to determine the trend in prevalence, awareness and control of hypertension in Malaysia.

**Methods:** Systematic searches were conducted in PubMed, Scopus, Ovid, Cumulative Index to Nursing and Allied Health Literature, Malaysian Medical Repository and Malaysia Citation Index published between 1980 and 2017. All original articles in English were included. Studies included were those on adults aged 18 years and above. Studies of prevalence in children and adolescents and pregnancy related hypertension were excluded. Two authors independently reviewed the studies, carried out data extraction and performed quality assessment. Heterogeneity between studies and publication bias was assessed and effect size was pooled by the random effect model.

**Results:** Fifty-six studies with a total of 241,796 subjects were included. The prevalence of hypertension throughout Malaysia varied (I^2^ = 99.3%). The overall pooled prevalence of hypertension over the past 4 decades was 28.2% in adults aged 18 years and older (95% CI: 26.1 – 33.3) and the prevalence in those 30 years and older was 40.0% (95% CI: 35.3-44.8).

For subgroup analysis, the prevalence of hypertension in male aged 18 and above was 31.4% (95% CI: 26.5 - 36.2) and 27.8% in female (95% CI: 20.7 – 34.9). The prevalence of hypertension among the ethnic groups aged 18 years and above were 37.3% in Malays (95% CI: 32.9 – 41.7); 36.4% in Chinese (95% CI 31.6 - 41.2) and 34.8% in Indians (95% CI: 31.2-38.4). The prevalence of hypertension was the lowest in the 1980s (16.2%, 95% CI: 13.4-19.0%), increases up to 36.8% in the 1990s (95% CI: 6.1-67.5), then came down to 28.7% (95% CI: 21.7-35.8) in the 2000s and 29.2% (95% CI: 24.0-34.4) in the 2010s. The prevalence of awareness was 38.7% (95% CI: 31.7 – 45.8) whereas the control of hypertension of those on treatment was 33.3% (95% CI: 28.4 – 38.2).

**Conclusion:** Three in 10 adults aged 18 years old and above have hypertension, whereas four in 10 adults aged 30 years old and above have hypertension. Four out of 10 are aware of their hypertension status and only one-third of them who were under treatment achieved control of their hypertension. Concerted efforts by policymakers and healthcare professionals to improve the awareness and control of hypertension should be of high priority.

## Introduction

Hypertension is a common public health problem over the past several decades [1-3]. It is one of the major risk factors for cardiovascular diseases (CVD) like stroke, heart failure and ischemic heart disease, [4]. Studies have been done repeatedly over the last 40 years on prevalence, awareness and control of hypertension in Malaysia. The National Health and Morbidity Surveys (NHMS), which are nationwide studies, were done every 10 years since 1986 in Malaysia. The NHMS has also shown the trend of hypertension in different genders and ethnicities. Besides the NHMS, several other studies also examined the prevalence of hypertension in specific groups and settings that were different from that of the NHMS [5, 6]. Prevalence in different settings may provide a slightly different picture and their accompanying set of problems. Because of the different settings and their conceivably different prevalence, there is a need for a systematic review and meta-analysis to gather and review all the information provided by these studies. Furthermore there is no systematic review that has been done to provide more information on the changes and trends of prevalence of hypertension in the past 4 decades in Malaysia as well. Hence, this systematic review aimed to determine the prevalence, control and awareness of hypertension in Malaysia over the past 4 decades.

## Methods

### Literature search strategy

Literature search was done between August 2017 and February 2018. Eligible articles were identified independently by two authors (CZY, TCH) through performing an initial screen of titles or abstracts, followed by full text review. Disagreements on study inclusion, data extraction and quality assessment were resolved by discussion between the 2 authors or by other senior authors. We comprehensively searched through 6 databases which are Pubmed, Ovid, Scopus, Cumulative Index of Nursing and Allied Health Literature (Cinahl), MyCite [7] and MyMedR [8]. The search strategy was based on terms related to: (prevalence) and (awareness) and (control) and (hypertension or high blood pressure) and (Malaysia) and combination of these using Boolean operators, adapted from previous reviews, and a combination of expanded MeSH term and free-text searches were conducted. A combination of expanded MeSH term and free-text searches were used as shown in Appendix 3. Cross reference of all selected articles were scanned for additional studies. If more than one article from a study was published, the article that provided the most updated data was selected. For studies that used the same set of data, only the study with the primary data was chosen, in view of possible duplication of data.

### Inclusion criteria

Studies conducted in Malaysia and published between 1 January 1980 and 31 December 2017 fulfilling the following criteria were included into the analysis: (1) Studies including the general population who are 18 years old and above; 2) studies on prevalence of hypertension diagnosed using digital automated or mercury sphygmomanometer 3) studies which were on either prevalence, awareness, control of blood pressure or hypertension individually or any combination of the three; (4) hypertension was defined as BP≥140/90 mmHg according to JNC 5; (5) studies on prevalence at BP screening campaigns 6) prevalence which was based either on physician-diagnosed hypertension or on antihypertensive medications; (7) awareness was defined as knowing one’s own hypertension status or having been diagnosed as hypertension previously; (8) control was defined as achieving target blood pressure of less than 140/90 mmHg; (9) studies in English only.

### Exclusion criteria

We excluded intervention studies, case studies, pharmacogenetic studies, case series which only included qualitative data, comments or letters, audits, narrative reviews, conference proceeding, opinion pieces, methodological, editorials, animal studies or any other publications lacking primary data and/or explicit method descriptions.

### Quality assessment and data extraction

Each article was undergone quality assessment by two authors using a modified critical appraisal checklist (Appendix 4) [57]. Each article’s quality was graded as ‘high quality’ if it scores ≥7/11; or graded as ‘low quality’ if it scores score <7/11 [5, 6]. The scoring result was shown in Appendix 3. Study characteristics (first author, year of publication, place and year of study, study setting, sampling method, sample size, blood pressure apparatus and classification cut-offs), participant characteristics (age group, gender, ethnicity and geographical origin), and prevalence (prevalence of hypertension, awareness and control) were extracted onto pre-coded spreadsheets independently by two authors (CZY, TCH). Data were extracted at the lowest possible disaggregate level referred to as subpopulation here. This review is presented according to the PRISMA guideline [10].

### Statistical Analysis

Sensitivity analyses were performed by discarding low quality studies, removing outlier subpopulations (point estimates > 3SD), or removing smaller subpopulations (size < 100). Effect size of interest was the proportion of individuals with hypertension. For estimating secular trend of prevalence, point estimates for four separate decades were calculated. We used OpenMetaAnalyst for data analyses [11]. Heterogeneity between studies for the pooled estimates was examined using I^2^ where an I^2^ of ≥75% suggests considerable heterogeneity. If the heterogeneity of the trials were ≥75%, a random effects model was used to pool the prevalence, awareness and control of hypertension. Subgroup analysis was done to examine the prevalence of hypertension by gender, age group, ethnicity, setting and geographical origin. Publication bias was assessed by visual inspection of funnel plots. Statistical significance was set at p value <0.05. The analyses was available in Appendix 4.

**Figure 1:**
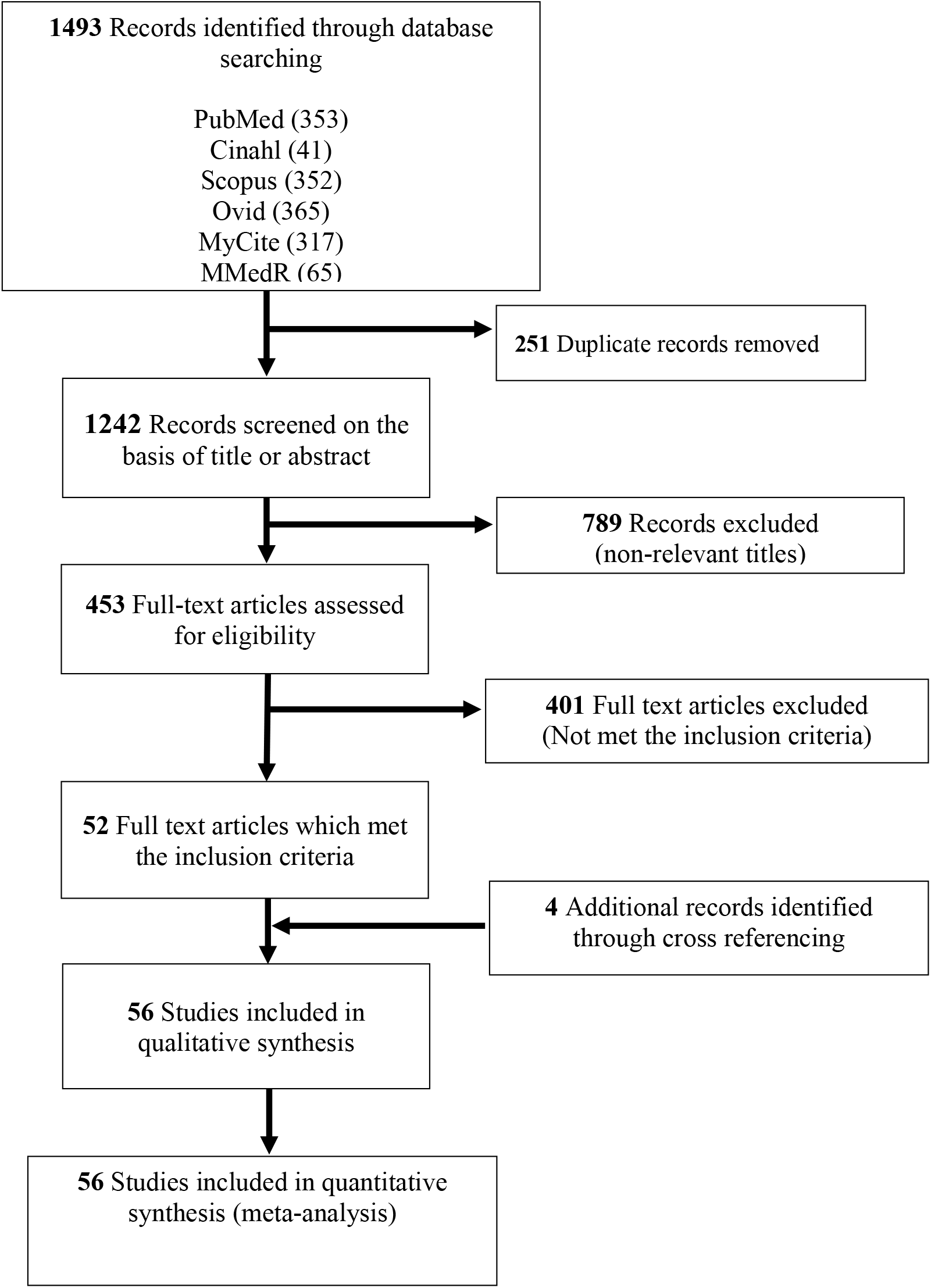
PRISMA flow diagram of the literature screening process. Numbers indicate the article count retained at each step of the process.

## Results

### Characteristics of studies included

We identified a total of 1493 manuscripts in the initial search as shown in Figure 1. 1242 studies were retrieved for further assessment after duplicated records were removed (n=251). Evaluation of the inclusion/exclusion criteria showed that 52 studies fulfilled eligibility, together with another 4 studies identified from cross-referencing making a total of 56 studies to be included into our systematic review. Using a modified critical appraisal checklist, quality assessment showed that 51 studies (50 community studies and 1 study done in healthcare setting) were of high quality (Appendix 3). A total sample size of 241,796 respondents from Malaysia were included into the analysis. Overall, the total subjects comprised of 51352 Malays, 22005 Chinese, 8819 Indians and 8638 other ethnicities. The setting of the study was examined in subgroup analysis; 51 studies were conducted in the community setting while the remaining 4 studies were conducted in hospital setting and 1 study in primary care clinic. The characteristics of each study was detailed in Appendix 1.

### Prevalence of hypertension

Table 1 shows the prevalence of hypertension in Malaysia according to different age groups, gender, ethnicities, setting, geographical origin, decades of study population and blood pressure measurement instruments used. The overall mean prevalence of hypertension in adults ≥ 18 years old and above was 29.7 % (95% CI: 26.1-33.3), while in those aged ≥ 30 years old, the prevalence was 30.8%, (95% CI: 25.5-36.2). Prevalence of hypertension was 16.2% (95% CI: 13.4-19.0) in the first decade (1980-1989); 36.8% (95% CI: 6.1-67.5) in the second decade (1990-1999); 28.7% (95% CI: 21.7-35.8) in the third decade (2000-2009) and 29.2% (95% CI: 24.0-34.4) in the fourth decade (2010-2017)

**Table 1:**
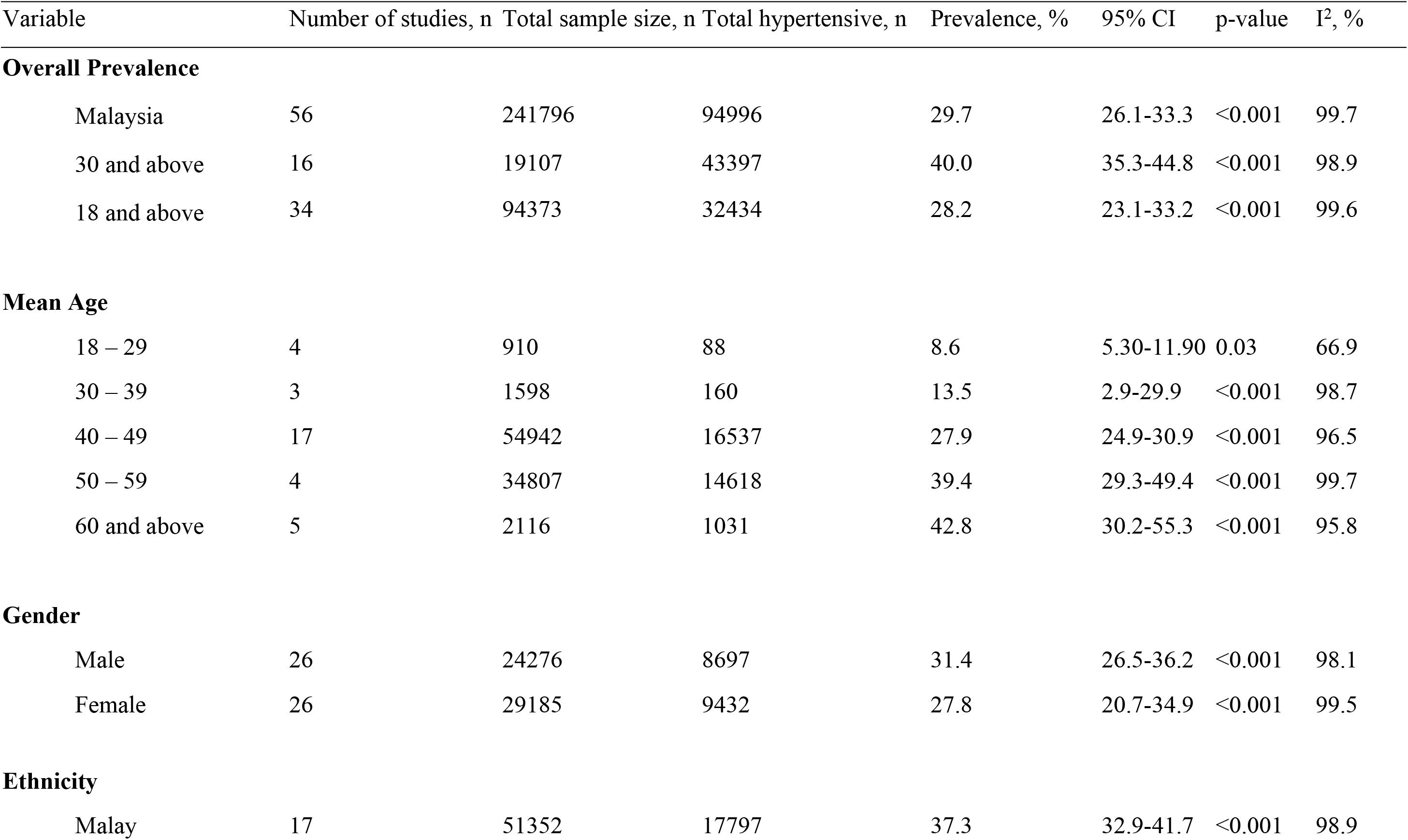
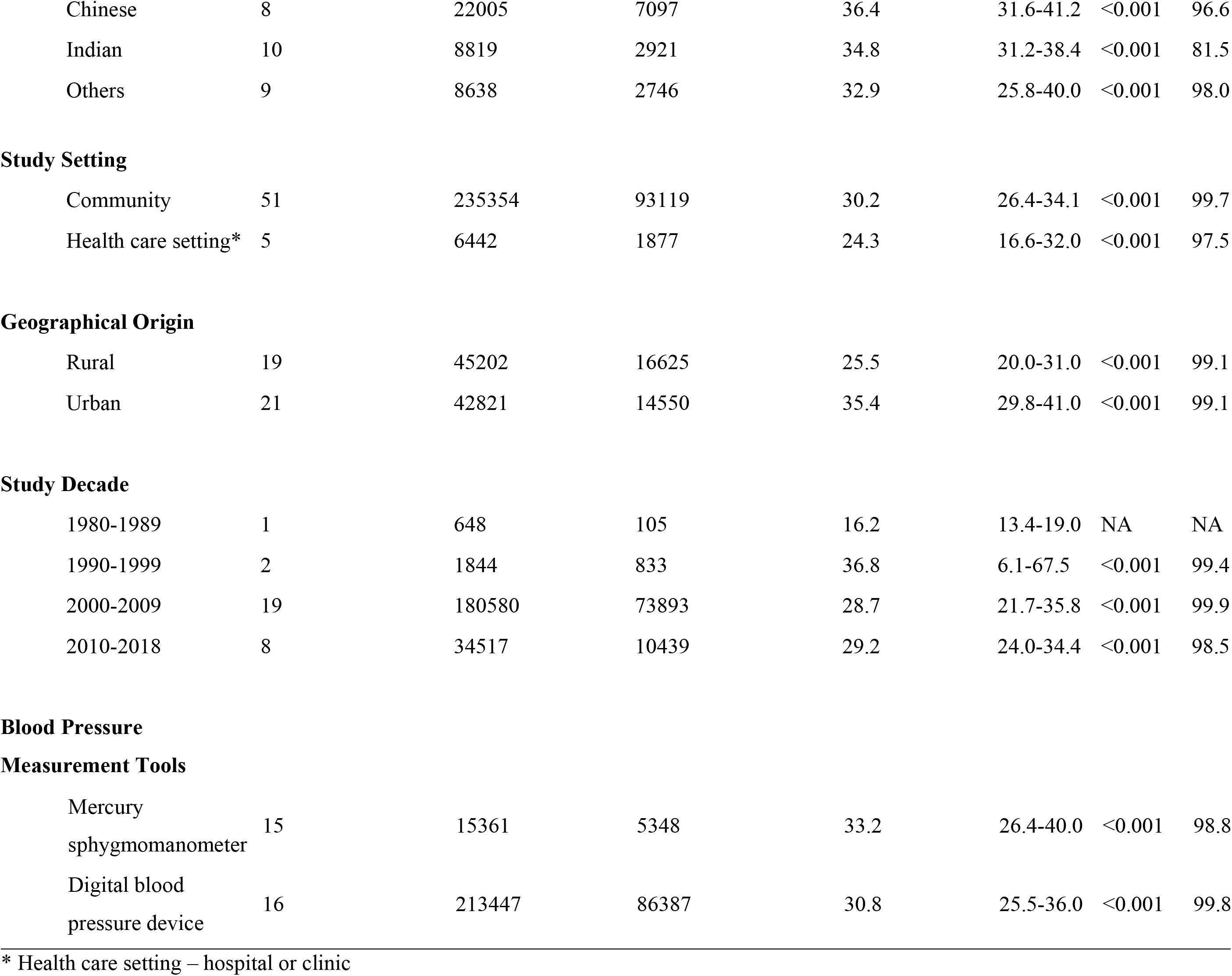
Pooled prevalence and 95% confidence interval of hypertension and its subgroup analysis

The prevalence of hypertension increases with age, from 8.6% among adults 18 to 29 years old to 42.8% among adults aged 60 years old and above. Among adults aged 18 years old and above, the prevalence of hypertension was seen to be higher in male compared to female (31.4%, 95% CI: 26.5-36.2 versus 27.8%, 95% CI: 20.7-34.9 respectively) in adults aged 18 years old and above. The prevalence of hypertension was also found to be highest among Malays (37.3%, 95% CI: 32.9-41.7), followed by Chinese (36.4%, 95% CI: 31.6-41.2) and Indian (34.8%, 95% CI: 31.2-38.4). Based on the study setting, the prevalence of hypertension was 30.2% (95% CI: 26.4-34.1) in communities and 24.3 (95% CI: 16.6-32.0) in health care setting. Prevalence of hypertension in rural areas was 25.5% (95% CI: 20.0-31.0) and 35.4% (95% CI: 29.8-41.0) in urban area. The prevalence of hypertension in studies using mercury sphygmomanometer was 33.2% (95% CI: 26.4-40.0) while those using digital blood pressure device was 30.8% (95% CI: 25.5-36.0).

### Awareness of hypertension

The overall awareness of hypertension in Malaysia as identified in 10 studies was 38.7% as shown in Table 2. Only 3 studies had examined the gender difference of awareness of hypertension. The awareness of hypertension among male hypertensive patients ranged from 49.5% to 92.0%, whereas, the awareness of hypertension among female hypertensive patients ranged from 56.4% to 71.0% [12, 54, 57]. In term of ethnicity, only 1 study examined the ethnic difference of hypertensive awareness, meanwhile there were 2 studies which their study populations were all Malays. The awareness of hypertension among Malays ranges from 28.0% to 55.8%, while the awareness of hypertension among Chinese, Indian and other minorities were 52.3%, 51.7% and 40.4% respectively. Based on their geographical origin, only 1 study examined the difference in awareness of hypertension, while there were 3 studies done in rural communities. The awareness of hypertension among hypertensive patients living in rural areas ranged from 28.0% to 52.35, while the awareness of hypertension among those who stay in urban areas was 54.1% based on a study done in 18 urban areas across all 7 states in Malaysia.

**Table 2:** Pooled Awareness, Pooled Control and 95% Confidence Interval of Total Hypertensive Patients and its Subgroup Analyses

| Variable | Number of studies, n | Total sample size, n | Total hypertensive, n | Prevalence, % | 95% CI | P-value | I<sup>2</sup>, % |
| --- | --- | --- | --- | --- | --- | --- | --- |
| **Overall Awareness** |  |  |  |  |  |  |  |
| Malaysia | 10 | 25794 | 11802 | 38.7 | 31.7-45.8 | <0.001 | 99.0 |
| **Overall Control** |  |  |  |  |  |  |  |
| Malaysia | 8 | 4698 | 1471 | 33.3 | 28.4-38.2 | <0.001 | 85.2 |
| **Gender** |  |  |  |  |  |  |  |
| Male | 3 | 559 | 180 | 37.1 | 26.0-48.2 | <0.001 | 59.1 |
| Female | 3 | 663 | 207 | 30.4 | 16.0-44.8 | <0.001 | 80.7 |
| **Ethnicity** |  |  |  |  |  |  |  |
| Malay | 2 | 1656 | 486 | 29.3 | 27.2-31.5 | 0.68 | 0 |
| Non-Malay | 3 | 2127 | 651 | 32.5 | 29.9-41.3 | 0.079 | 52.2 |
| **Geographical Origin** |  |  |  |  |  |  |  |
| Rural | 3 | 1147 | 317 | 34.1 | 15.5-52.7 | <0.001 | 94.7 |
| Urban | 1 | 1116 | 407 | 36.5 | 33.6-39.3 | NA | NA |

### Control of hypertension

Among the hypertensive subjects who were aware of being hypertensive, 33.3%, 95%CI: 28.4% to 38.2%) achieved blood pressure control. The heterogeneity between the studies was significant (I^2^ = 85.2%, p<0.0001). Men had a slightly better control than women in our analysis (37.1% vs 30.4% respectively). In examining the ethnic groups, it was found that Malay has a control rate of blood pressure at 29.3%, while non-Malay had a control rate of 32.5%. The control of hypertension was higher in urban dwellers than in the rural area (36.5% vs 34.1% respectively).

### Sensitivity analyses

The main analysis for prevalence of hypertension was rerun by removing one subpopulation at a time. The pooled estimates did not vary much from the original analysis during each removal. Removal of five low quality studies or the smaller (size < 100) subpopulation did not affect the original estimate. (Appendix 6)

## Discussion

To the best of our knowledge, this systematic review is the first in Malaysia to describe the prevalence and its trends over 4 decades, awareness and control of Hypertension. Multiethnicity is one of the special features in Malaysia that would cause variation in prevalence, awareness and control of hypertension.

### Prevalence

The overall prevalence of hypertension in those aged 18 years and older in our study was 29.7% (95% CI 26.1 – 33.3). The overall prevalence of hypertension in Malaysia is within the range of worldwide prevalence of hypertension (20% to 50%) as described in the systematic review by Kearney et. al. [68]. Our study shows that Malaysia’s prevalence of hypertension is within the worldwide range. In comparing other Asian countries, the prevalence of hypertension in Malaysia is higher than Thailand (24.7%), Singapore (23.5%) and China (25.2%) [69, 70, 71]. A review showed that this prevalence was as high as developed countries despite being a developing country [72]. In fact our prevalence of hypertension is even higher than that of United States by 0.7% [73].

The heterogeneity of the overall prevalence in this study was high. This could be due to our wide variation of sample size in our review, which ranged from 56 to 106527. Sample size is one of the most important factors to determine the prevalence of hypertension as a previous study had demonstrated a significant association between sample size and the prevalence of hypertension [74]. Besides sample size, the wide variation in prevalence found in our study could be due to the different settings of data collection. For example, data collected in community settings may differ from the data collected from hospitals or primary health care clinics as the visitors of health care settings are more likely to have hypertension and other underlying comorbidities which tend to increase the prevalence. However, surprisingly in our study, the prevalence of hypertension was lower in healthcare settings compared to community settings. The possible reason could be due to the study population in those healthcare settings had low cardiovascular risk profile; out of 5 studies, 3 studies were done among healthcare workers without any comorbidities [16, 32, 47] and 1 study was only done in non-hypertensive patients by excluding those who were known hypertensive patients [41]. Furthermore, the mean age of the study population in studies done in healthcare setting is relatively younger, which ranges from 43.5 years old to 49.4 years old, compared to those studies done in communities [16, 32, 30, 41, 47].

Geographical origin also played an important role in causing such high heterogeneity in our study. In our study, we had a 9.9% difference in both geographical origin, in which urban dwellers had a higher prevalence of hypertension compared to those living in rural areas. This could be due to a more sedentary lifestyle and higher prevalence of obesity seen in urban dwellers and these themselves are risk factors for developing hypertension. A rural population is more likely to be in occupations or physical activities that require more energy expenditure [75].

It has been reported that digital blood pressure monitor show poor agreement with mercury sphygmomanometer in measuring blood pressure, thus influencing the difference in prevalence of hypertension when different instruments were used [76]. Despite showing a slight higher prevalence of hypertension using mercury sphygmomanometer compared to digital blood pressure device in our review, the difference between the two devices was not substantial. Furthermore, in view of the safety issues of possible mercury leakage, the difference between the prevalence of hypertension using these two devices should not be taken into consideration during our daily clinical practice. As such, it is reasonable to use digital blood pressure device as it is safer and more convenient than the “gold standard” mercury sphygmomanometer.

### Trend of hypertension

Table 1 shows the increasing trend of prevalence of hypertension for over the past 30 years since the 1980s. What is striking in our study is the low prevalence of hypertension in 1980s. We found that only one study describing the prevalence of hypertension in the 1980s. This study was done among the Kadazan and Bajau ethnic group, a minority group in a rural part of Sabah [28]. Hence it is not surprising that the prevalence then was so low as not only was this a rural population but it was the era before urbanisation, unhealthy lifestyle and increase in economic wealth took over Malaysia. Otherwise, the prevalence of hypertension was found to be highest in the decade of 1990s with the prevalence of 36.8% (95% CI 6.1 – 67.5). This gradually came down to 28.7% in 2000s (95% CI 21.7 – 35.8) and 29.2% in 2000s (95% CI 24.0 – 34.4). On the other hand while the trend in US was one of declining prevalence, it was increasing in Malaysia. This trend was similar with the trend of hypertension found in United States population from 1980s to 2010s according to NHANES study, which was around 30.0% [30, 73, 77]. Besides that, it was surprising to have such a stand-out prevalence in the 1990s. This was due to the fact that among all 30 studies that specified their study date, only 2 studies that was conducted in the 1990s. One study, which had a prevalence of hypertension at 21.1%, was done in three rural communities in Bagan Datoh with a wide variation of citizens from different age group [52], whereas another study was done in another three semi-rural areas in Kuala Langat which its study respondents are coming from older age groups, ranging from 55 years old to 95 years old with a mean age of 65.4 years old [25]. This significantly increases the overall pooled prevalence of hypertension if we only took these 2 studies with extreme ends of prevalence into account.

### Age and Hypertension

Epidemiological studies have already shown that the prevalence of hypertension increases with age. Our review shows the same trend. Importantly, what was also seen in our study was that there was a doubling in the prevalence of hypertension in those aged between 40 to 49 years old (27.9%) while it was only 13.5% in those aged 30-39 years old. While comparing our result to developed country, we also found a similar doubling phenomenon in the prevalence of hypertension, but only later in age, which is 63.1% in age group 60 years old and above from 33.2% among those from age group 40-59 years old [79]. This showed that the prevalence of hypertension in Malaysia starts at an earlier age compared to the developed country which makes young hypertension to be an anticipating issue to be concerned in the future in Malaysia. This is expected as older age is closely related with hypertension because of the alteration in the arterial structure and ongoing calcification that leads to increased arterial stiffness [78]. In short, with increasing age it is expected that the prevalence of hypertension will rise. However, focussing on the older population aged 60 years old and above, the prevalence of hypertension in this age group in Malaysia is found to be the lowest among Asian countries such as Singapore (73.9%), Korea (68.7%), India and Bangladesh (65%), Taiwan (60.4%), Thailand (51.5%) and China (48.8%) [80, 81, 81, 83, 84, 85]. However this could be due to the fact that studies in Malaysia have been defining elderly as individuals aged 60 years old and above compared to other studies as mentioned earlier which defined them as individuals aged 60 years old and above. Hypertension among younger age group in Malaysia was found to be lower as well where 8.6% of study population 18 to 29 years old and 13.5% of study population 30 to 39 years old had hypertension. This is higher as compared to the developed country such as United States (7.3%) but almost similar to China (8.9% - 14%) and lower compared to India (17.72%) [86, 87, 88].

### Gender and Hypertension

The prevalence of hypertension among males was higher than females in Malaysia. This finding is similar to the National Health and Nutrition Examination Survey (NHANES) done in United States in put year here, which reported that men regardless of race and ethnicity had a higher prevalence of hypertension than women in the age group of 20 to 40 [86]. The gender differences in hypertension are due to both biological and behavioural factors [89]. From the biological aspect, the female sex hormone, oestrogen serves as a protective factor against hypertension and other cardiovascular diseases in women [90, 91, 92]. Meanwhile from the behavioural aspects, smoking prevalence is higher among males compared to females [90, 91, 92, 93] and cigarette smoking is a known risk factor for hypertension [94, 95]. This explains why the prevalence of hypertension is higher in males.

### Awareness

In our review, 38.7% (95% CI 31.7-45.80, I^2^ 98.95) were aware of their hypertension status. This finding is lower than that in United States (63%) [77], Singapore (69.7%) [80] and Korea (91.7%) [81]. On the other hand, the awareness of hypertension in Malaysia is higher than that of India (25.1%) [82] and Indonesia (35.8%) [69]. Our level of awareness is still poor as it means that almost 2 put of 3 adults with hypertension go undetected and untreated. Unless these individuals are detected, they will continue to contribute to the high cardiovascular mortality and morbidity. Steps that can be done to improve awareness could include some of the initiatives such as the May Measurement Month 2017 which is nationwide blood screening program which is done by a joined cooperation of International Society of Hypertension and University Malay Medical Center in conjunction with World Hypertension Day. This nationwide screening of adults aged 18 years old and above was conducted through various kind of events such as health campaigns at clinics, hospitals and community centres, family day events as well as charity runs from 1 April 2017 to 31 May 2017 in 42 different centres [96].

As for the gender difference of awareness of hypertension, despite one study had a perceivably large sample size of study population done across 7 states of Malaysia [12], the other two studies were done in two conceivably different settings with relatively higher awareness than the general population as reported in Ab Majid et al., 2018 [97]. One study was done in a residential home with high ratio of caregiver to residents in the home, which will certainly increase the awareness of the residents on hypertension [54]. Another study was done in university staff with high educational level which certainly increases the prevalence of awareness of hypertension [57]. In term of ethnicity, only one study examined the ethnic difference of awareness of hypertension [12], while in other two studies, their study population were all elderly Malays [58] and Malay villagers in a rural communities respectively [66]. Same goes to geographical origin, in which only one study examined the difference in awareness [12] while the other three studies were all focussed in rural communities rather than examining the geographical difference of hypertensive awareness [42, 48, 58]. With these in mind, it is safe to assume that there will be much biases and high heterogeneity, and therefore pooled analyses were not done for all these subgroups.

### Control

The control of hypertension in Malaysia was 33.3% (29.1-39.2) and this is much lower than that in developed countries such as United States (53%) [13]. On the other hand, this figure is higher than nearby countries like China (13.8%) [69, 84], Hong Kong (25.8%) [69, 98] and Philippines (27.0%) [69, 99]. This could be due to the fact that Malaysia has been improving health care facilities with more clinics and hospitals being built ensuring better access to health care [100]. Males achieved a better control of blood pressure than females. Despite being consistent with local nationwide study, this is surprising as females are more likely to have better health-seeking behaviour, including the fact that female is not the risk factor of hypertension [93]. Urban dwellers had better control of blood pressure, which correlates with a study done in Southern China which showed the similar result [9]. This may be due to limited access to the health care facilities in the rural areas despite the increasing number of rural clinics throughout the past 4 decades in Malaysia [100]. It seems likely that the poorer health awareness among rural dwellers or lower socioeconomic profile still remains as an important barrier to their visiting health care facilities and receiving proper treatment.

### Strengths and Limitation

The main advantage of this review was the large sample size which resulted in the highly precise pooled estimate. Besides that, this review showed the trend of hypertension throughout the past 4 decades in Malaysia. There were certain limitations in this study. Firstly, there are many studies which did not report the prevalence of hypertension according to gender, ethnicity and geographical origin. Secondly, there are also many unpublished data or grey literature which were not included in this study. Thirdly, estimates for earlier time periods were based on fewer studies when compared to estimates for later periods, which may have resulted in less precise result.

### Suggestion for future research

Future studies on prevalence of hypertension can be improved by addressing some of these issues. The prevalence of hypertension according to gender, ethnicity and geographical origin should be studied in more detail. Studies also should avoid using non-random sampling method as it would lead to bias in the study. Besides that, future studies should also emphasize on adequate or larger sample size which are more representative of a population.

## Conclusion

In conclusion, one-third of Malaysian adults are hypertensive. Prevalence of hypertension is higher in urban dwellers than those in rural areas. Unfortunately, only about one third of the hypertensive patients were aware of their hypertension status and only one third of the hypertensive patients achieved target blood pressure control. In view of these findings, urgent steps to improve health promotion and health education have to be made on a larger scale. Although this review showed a decreasing trend of prevalence of hypertension throughout the past 4 decades, nonetheless the awareness and control of blood pressure among Malaysians has yet to be improved.

## Authors’ contributions

Conceived and designed the study: CSM CYC NKD HFK CZY TCH. Performed the study: CZY TCH LKW. Analysed the data: CZY SMJ TCH LKW. Wrote the paper: CZY TCH. Data interpretation: CZY SMJ CSM TCH CAT NKD. Critical revision to the manuscript: CYC CSM CAT NKD HS VD.

## Abbreviation

I: Heterogeneity
CI: Confidence Interval
CVD: Cardiovascular Diseases
NHMS: National Health and Morbidity Survey
Cinahl: Cumulative Index to Nursing and Allied Health Literature
MyCite: Malaysian Citation Index
MyMedR: Malaysian Medical Repository
MeSH: Medical Subject Headings
JNC: Joint National Committee
PRISMA: Preferred Reporting Items for Systematic Reviews and Meta-Analyses
SD: Standard Deviation
NHANES: National Health and Nutrition Examination Survey

